# The Mechanism underlying B cell Developmental Dysfunction in Kawasaki Disease Based on Single-cell Transcriptomic Sequencing

**DOI:** 10.1101/2023.02.21.529340

**Authors:** Qiuping Lin, Zhen Wang, Guohui Ding, Guang Li, Liqin Chen, Qingzhu Qiu, Sirui Song, Wei Liu, Xunwei Jiang, Min Huang, Libing Shen, Tingting Xiao, Lijiang Xie

## Abstract

**Background:** Kawasaki disease (KD) is an acute systemic vasculitis that can lead to acquired heart disease in children mostly from in developed countries. The previous research showed that B cells in KD patients underwent a profound change in both the cell numbers and types after intravenous immunoglobulin (IVIG) therapy.

**Methods:** We performed the single-cell RNA-sequencing for the peripheral blood mononuclear cells (PBMCs) from three febrile patients and three KD patients to investigate the possible mechanism underlying B cell developmental dysfunction in KD. A previously published single-cell sequencing KD dataset (GSE168732) was also utilized in study for sample size expansion and validation. The comprehensive single-cell data analyses were applied for our dataset and GSE168732 dataset including single-cell trajectory analysis. To validate the immune disorders in KD, we measured immune-related indicators from 28 KD and 28 febrile patients.

**Result:** Overall single-cell expression profiles show that the biological processes of immunity, B cell activation pathway and their related biological entities are repressed in KD patients before IVIG treatment compared to febrile patient and KD patients after IVIG treatment. The differentially expressed gene analyses further demonstrate that B cell signaling pathway is downregulated in B cells and plasma blast cells of KD patients before treatment while cell cycle genes and MYC gene are upregulated in dendritic cells (DCs) and hematopoietic stem and progenitor cells (HSPCs) of KD patients before treatment. The biological process of immune response is upregulated in the HSPCs of KD patients before treatment in our dataset while the biological process of inflammatory response is upregulated in the HSPCs of KD patients before treatment in GSE168732 dataset. Single-cell trajectory analyses demonstrate that KD patients before treatment have a shortened developmental path in which B cells and T cells are failed to differentiate into separate lineages. HSPD1 and HSPE1 genes show an elevated expression level in the early cell development stage of KD patients before treatment accompanied with the repression of MYC, SPI1, MT2A and UBE2C genes. Our analyses of all B cells from KD patients before treatment show most of B cells are arrested in a transitional state with an ill developmental path compared with febrile patients and KD patients after treatment. The percentage and absolute value of CD8 T cells in KD were lower than those in febrile patients. The ratio of CD4/CD8 in KD was higher than it in febrile patients. The serum levels of IgG and IgM in KD were lower than those in febrile patients.

**Conclusions:** Our results indicate that the immune premature HSPCs accompanied with the abnormal expression dynamics of cell cycle and SPI1 genes are the mechanism underlying B cell developmental dysfunction in KD patients.

**Funding:** This work is jointly supported by National Natural Science Foundation of China (82170518) and the Shanghai Science and Technology Committee research Funding (22Y11909700) and Shanghai Jinshan District medical key specialty Funding (JSZK2023A04).

## Introduction

Kawasaki disease (KD) is an acute systemic vasculitis that occurs predominantly in children under 5 years of age, which is the most common cause of acquired heart disease in children in developed countries (1). Coronary artery lesion (CAL) is a major complication of KD. High-dose intravenous immunoglobulin (IVIG) combined with aspirin is the standard treatment for KD (2). The diagnosis of KD mainly depends on clinical features. Several febrile diseases have similar clinical manifestations to KD, such as scarlet fever, EB virus infection, juvenile idiopathic arthritis, measles and adenovirus infection, leading to difficulties in the diagnosis and early treatment of KD (3). The etiology of KD so far remains unknown. The widely proposed theories have been in the categories of environmental fungal toxin exposure (4), autoimmune pathogenesis, and infectious diseases, but none of them have been proven (5). The most widely accepted etiologic hypothesis suggested that KD was caused by an infectious agent, which infected children with a genetic susceptibility (6).

In recent years, the hygiene hypothesis, initially proposed to explain the etiology of allergic diseases, had been introduced to explain the pathogenesis of KD (7). The hygiene hypothesis assumes that lacking of early-life microbial exposure results in an impaired or delayed immune maturation, leading to an abnormal response to the microbes harmless to healthy population, thus triggering KD (8, 9).The KD incidence gradually increased after its emergence in 1967 (10). The epidemic data showed an important characteristic of KD that the disease is relatively rare in children less than 6 months or more than 5 years old, with a peak incidence in children between 6 and 24 months old in different ethnic groups (11–13). KD was followed by a prolonged period of antigen-specific ‘split’ T-cell anergy, which reflected a maturational defect in immune responsiveness of KD patients (14). It suggests that KD may be caused by the immune maturation delay in early childhood, which is probably due to a nearly perfect sanitary environment. The B cell development dysregulation may be the potential mechanism behind the immune maturation delay of KD (15). Genetic studies further showed that KD susceptibility genes are involved in B cell development and function (16–18), underscoring the crucial role of B cell immunity in KD. Though these evidences related the hygiene hypothesis of KD to B cell development, the possible pathogenic mechanism has not been investigated at the cellular level.

Single-cell transcriptomic sequencing (scRNA-seq) is a relatively new tool for quantifying gene expression profile for an individual cell. Recent studies had utilized scRNA-seq data to study the immune disorders in KD (19–21). They revealed the dynamic changes of immune cells, differential gene expression and even biological processes in KD compared with other conditions, providing new insights into the etiology of KD and the therapeutic mechanism of IVIG. In this study, we obtained the scRNA-seq of peripheral blood mononuclear cells (PBMCs) from febrile patients, KD patients before and after IVIG treatment to explore the possible mechanism of B cell development dysfunction in KD. We found the biological processes related to immunity and B cell activation pathway were upregulated in the overall single-cell expression feature of febrile and KD patients after IVIG treatment, and the activation of cell cycle pathways was presented in dendritic cells (DCs) and hematopoietic stem and progenitor cells (HSPCs) pre-treatment KD. The analysis of peripheral blood hematopoietic stem and progenitor cells (HSPCs) confirmed the early immune activation propensity in pre-treatment KD. Single-cell trajectory analyses further revealed KD patients before treatment had a shorten cell development route and an ill cell differentiation outcome. The pseudo-time analyses indicated that a large proportion of B cells in KD patients before treatment are arrested in transitional state. Our study indicates that the abnormal expression pattern of HSPD1 and HSPE1 in immune premature HSPCs would result in the dysregulated B cell development and IVIG could rescue this process. Our study illustrates the probable etiology of KD and thus provides new insights into the diagnosis and treatment of KD.

## Result

### Clinical features and laboratory data of KD and febrile

In this study, we collected six peripheral blood samples from three KD patients (before/after IVIG treatment). All KD patients were sensitive to IVIG and did not develop CAL. We also collected peripheral blood samples from three febrile patients, which included two pneumonia patients and one pharyngitis patient. **Table 1** summarizes the clinical and laboratory information of KD and febrile patients.

**Table 1.**
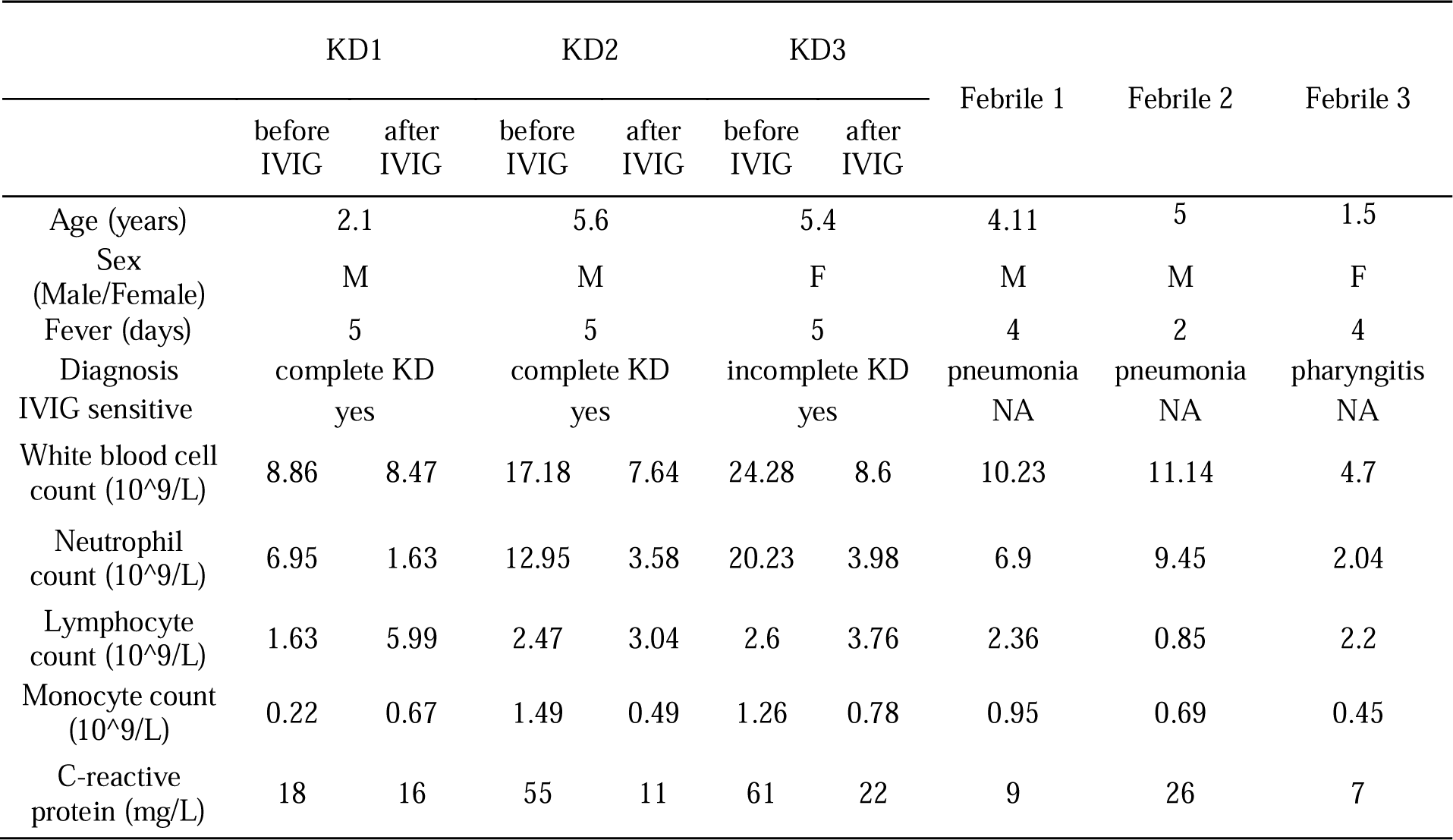
Clinical features and laboratory data for of KD and febrile patients.

|  | KD1 |  | KD2 |  | KD3 |  | Febrile 1 | Febrile 2 | Febrile 3 |
| --- | --- | --- | --- | --- | --- | --- | --- | --- | --- |
|  | before<br>IVIG | after<br>IVIG | before<br>IVIG | after<br>IVIG | before<br>IVIG | after<br>IVIG |  |  |  |
| Age (years) | 2.1 |  | 5.6 |  | 5.4 |  | 4.11 | 5 | 1.5 |
| Sex<br>(Male/Female) | M |  | M |  | F |  | M | M | F |
| Fever (days) | 5 |  | 5 |  | 5 |  | 4 | 2 | 4 |
| Diagnosis | complete KD |  | complete KD |  | incomplete KD |  | pneumonia | pneumonia | pharyngitis |
| IVIG sensitive | yes |  | yes |  | yes |  | NA | NA | NA |
| White blood cell<br>count (10<sup>9</sup>/L) | 8.86 | 8.47 | 17.18 | 7.64 | 24.28 | 8.6 | 10.23 | 11.14 | 4.7 |
| Neutrophil<br>count (10<sup>9</sup>/L) | 6.95 | 1.63 | 12.95 | 3.58 | 20.23 | 3.98 | 6.9 | 9.45 | 2.04 |
| Lymphocyte<br>count (10<sup>9</sup>/L) | 1.63 | 5.99 | 2.47 | 3.04 | 2.6 | 3.76 | 2.36 | 0.85 | 2.2 |
| Monocyte count<br>(10<sup>9</sup>/L) | 0.22 | 0.67 | 1.49 | 0.49 | 1.26 | 0.78 | 0.95 | 0.69 | 0.45 |
| C-reactive<br>protein (mg/L) | 18 | 16 | 55 | 11 | 61 | 22 | 9 | 26 | 7 |

### Immunological parameters in KD and febrile

To validate the immune disorders in KD patients, we measured immune-related indicators in 28 KD and 28 febrile patients (**Table 2**). There were 15 males and 13 females in both KD group and febrile group. There was no difference in age between KD and febrile (28.36±17.64 vs 28.82±10.89 months, p>0.05). There was no difference between fever days in KD and febrile (5.00±0.00 vs 4.82±0.98 days, p>0.05). The percentage (17.41±4.88% vs 23.45±6.51%) and absolute value (0.55±0.35 10^9/L vs 0.74±0.36 10^9/L) of CD8+T cells in KD were significantly lower than those in febrile (p<0.05). The ratio of CD4/CD8 ( 2.43±1.28 vs 1.59±0.69) in KD was significantly higher than it in febrile (p<0.05). The percentages and absolute values of CD3+T cells, CD4+T cells, CD16+56+/CD3-cells, and CD19+ cells in the two groups showed no differences (p>0.05). We also measured the serum levels of IgG, IgM, IgA, and IgE in KD and febrile. The levels of IgG (6.29±1.86 g/L vs 8.59±2.24 g/L) and IgM (0.91±0.34 g/L vs 1.14±0.34 g/L) in KD were significantly lower than those in febrile (p<0.05), while the levels of IgA and IgE in KD and febrile showed no difference (p>0.05).

**Table 2.** Immune-related indicators in KD and febrile patients.

|  | KD group ( n=28 ) | Febrile group ( n=28 ) |
| --- | --- | --- |
| CD3+(%) | 59.13±9.57 | 61.45±9.92 |
| CD8+(%) | 17.41±4.88* | 23.45±6.51* |
| CD4+(%) | 37.85±7.36 | 33.99±9.8 |
| CD16+56+/CD3-(%) | 8.73±5.56 | 8.55±5.46 |
| CD19+(%) | 30.99±8.09 | 28.17±8.55 |
| CD4+/CD8+(%) | 2.43±1.28* | 1.59±0.69* |
| CD3+(absolute count)(10<sup>9</sup>/L) | 1.86±0.93 | 2.13±0.17 |
| CD8+(absolute count)(10<sup>9</sup>/L) | 0.55±0.35* | 0.74±0.36* |
| CD4+(absolute count)(10<sup>9</sup>/L) | 1.18±0.64 | 1.17±0.78 |
| Lymphocytes+(absolute count)(10<sup>9</sup>/L) | 3.08±1.40 | 3.39±1.76 |
| CD16+56+/CD3-(absolute count)(10<sup>9</sup>/L) | 0.27±0.20 | 0.31±0.31 |
| CD19+(absolute count)(10<sup>9</sup>/L) | 0.94±0.44 | 0.95±0.48 |
| IgG ( g/L ) | 6.29±1.86* | 8.59±2.24* |
| IgA ( g/L ) | 0.71±0.49 | 0.73±0.36 |
| IgM ( g/L ) | 0.91±0.34* | 1.14±0.34* |
| IgE ( IU/ml ) | 264.83±622.88 | 263.27±325.62 |

\* represents a statistically significant difference ( P<0.05 )

### Single-cell transcription profiling of PBMCs in febrile and KD patients before/after treatment

To describe the general features of single-cell expression features of febrile and KD patients before/after IVIG treatment, we collected the PBMCs from 3 febrile patients (used as control), 3 KD patients before and after IVIG treatment. We also used a set of published KD data (GSE168732) to compare with our single cell sequencing result (21). After quality control and filter, the total number of detected cells was 69300, including 23072 cells from febrile patients, 19725 cells from KD patients before IVIG treatment (KD_BT), and 26503 cells from KD patients after IVIG treatment (KD_AT); the total number of detected cells from KD dataset of GSE168732 was 39114, including 16003 cells from KD patients before IVIG treatment (KD_BT_GSE168732) and 23111 cells from KD patients after IVIG treatment (KD_AT_GSE168732). To compare the scRNA-seq profiles between control and KD patents, we first integrated nine of our samples together and clustered the cells across samples according to their expression features (Figure 1A and Supplemental Figure 1A). We also integrated our febrile samples with six GSE168732 KD samples according to their expression features (Figure 1B and Supplemental Figure 1C). The detected cells could be classified into 12 major cell types covering B cells, CD4 T cells, CD8 T cells, CD14 monocytes (CD14 mono), CD16 monocytes (CD16 mono), dendritic cells (DC), erythrocytes (Eryth), gamma-delta cells (gdT), hematopoietic stem and progenitor cells (HSPCs), natural killer cells (NK), plasma blast cells (Plasmablast), and platelets. The canonical gene markers were used to validate major cell types (Supplemental Figure 1B and 1D). Figure 1c shows the percentage of major cell types for febrile patients and KD patients before and after IVIG treatment, as well as the KD samples from GSE168732 on the individual level. The scRNA-seq data demonstrated that KD patients after treatment had an increased percentage of CD8 T cells (p<0.05) and CD4 T cells (p<0.05) and a decreased percentage of monocytes (p<0.05) compared with those before treatment, which were well established by previous studies(21–23). Moreover, the number of B cells also shows a quite variance among febrile and KD patients before/after treatment. In both our dataset and GSE168732 dataset, there is a trend of more plasma blast cells in KD patients after treatment than KD patients before treatment (Figure 1A and 2B).

**Figure 1.**
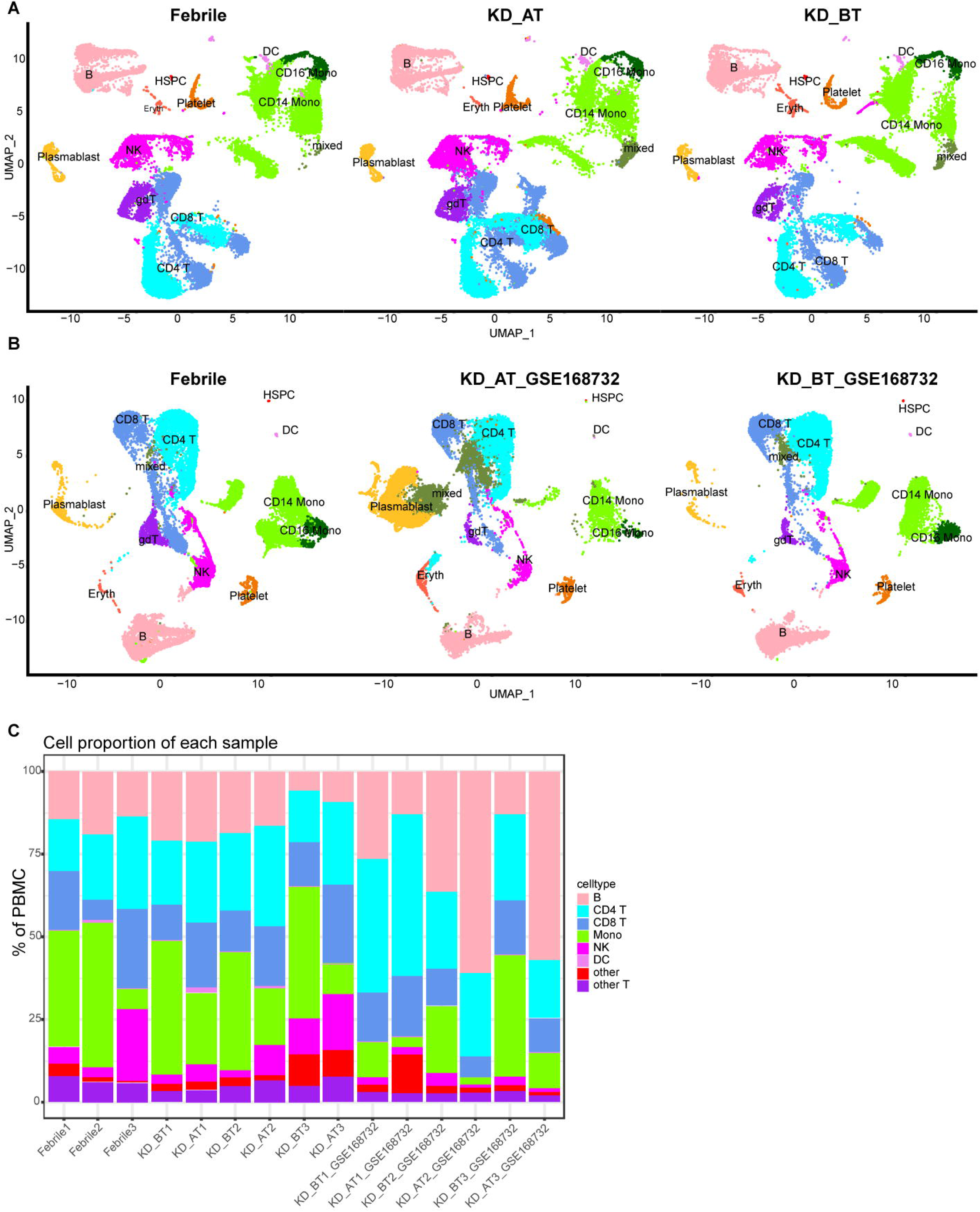
Single-cell profiling of PBMCs in our dataset and GSE168732 dataset. A. The integration single-cell profiling analysis of nine samples in our dataset which are 3 febrile patients (Febrile), 3 KD patients before treatment (KD_BT), and 3 KD patients after treatment (KD_AT). b. The integration single-cell profiling analysis of three samples in our dataset and six samples in GSE168732 dataset, which are 3 febrile patients (Febrile), 3 KD patients before treatment (KD_BT_ GSE168732), and 3 KD patients after treatment (KD_AT_ GSE168732). c. The proportion of different cell types in each individual sample. The inferred cell types are marked with different colors.

**Figure 2.**
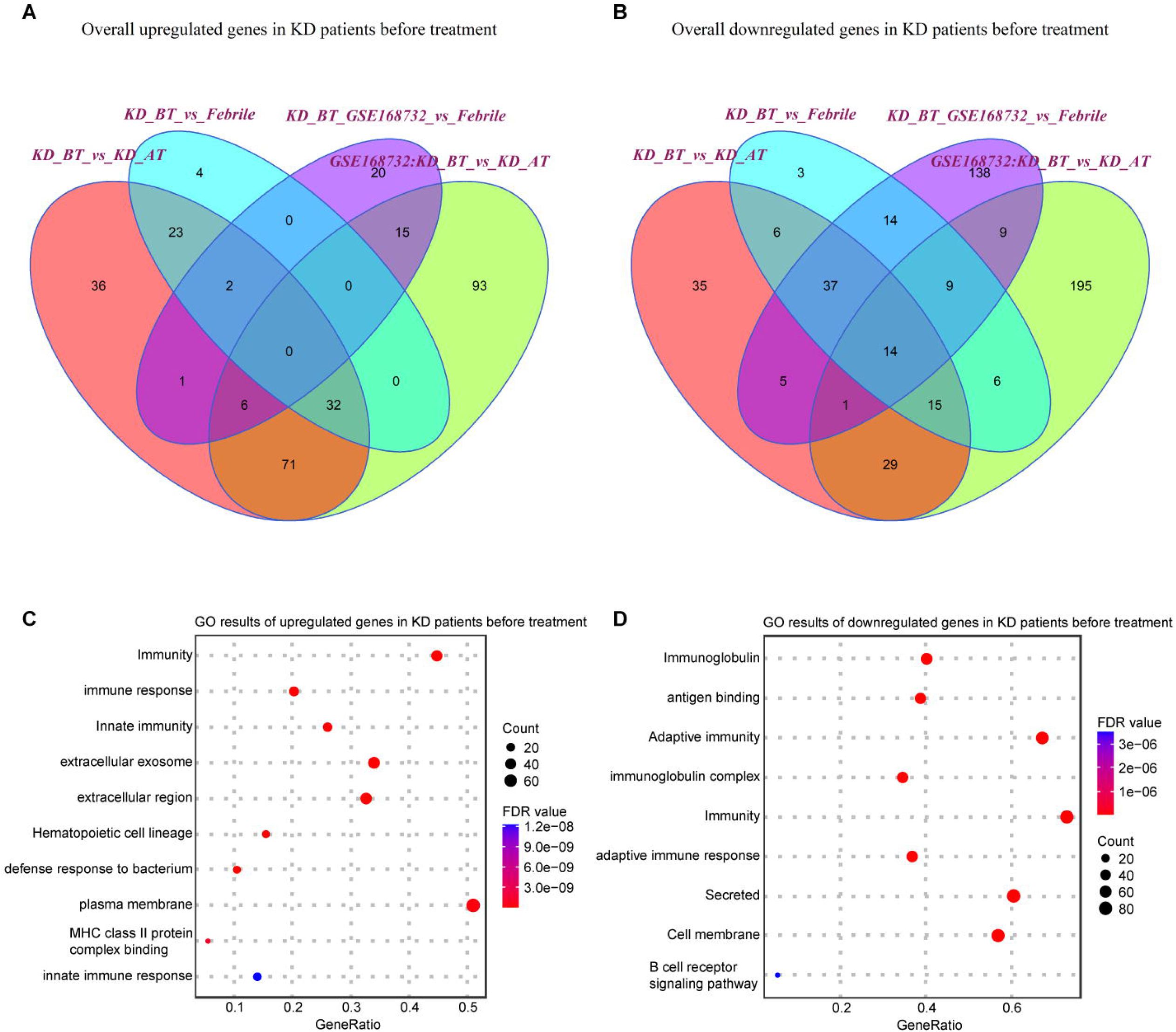
Expression analyses of all single-cells in our dataset and GSE168732 dataset. A. Venn diagram of upregulated genes in all single cells for KD patients before treatment in our dataset and GSE168732 dataset. B. Venn diagram of downregulated genes in all single cells for KD patients before treatment in our dataset and GSE168732 dataset. C. GO term enrichment analysis of 71 common upregulated genes in all single cells for KD patients before treatment. D. GO term enrichment analysis of 29 common downregulated genes in all single cells for KD patients before treatment.

### Overall expression features of all single cells in febrile and KD patients before/after treatment

We first examined the expression features of all single cells in febrile and KD patients before/after treatment. The differentially expressed genes (DEGs) were identified in KD patients before treatment using febrile and KD patients after treatment as background. There are 303 upregulated genes and 516 downregulated genes (p<0.05) detected in KD patients before treatment from our dataset and GSE168732 dataset (Figure 2A and 2B). 112 out of 303 upregulated genes and 130 out of 516 downregulated genes are shared by both datasets. Next, we performed gene ontology (GO) analysis on these shared upregulated and downregulated genes (Figure 2C and 2D). There is a common biological process of immunity found in shared upregulated and downregulated genes, although the genes behind this biological process are totally different in KD patients before treatment. The shared upregulated genes in KD patients before treatment are mainly involved in innate immunity and extracellular region while the shared downregulate genes in KD patients before treatment are mainly involved in immunoglobulin, adaptive immunity, and cell membrane. It shows that KD patients before treatment and febrile patients/KD patients after treatment rely on different immune mechanisms. The latter is more inclined to utilize secreted immunoglobulin as the measure of adaptive immune response. Notably, we found that B cell receptor signaling pathway are downregulated in KD patients before treatment, which proposes B cell related pathology for KD patients before treatment and is consistent with the previous study (21).

### Distinct expression features of B cells, plasma blast cells, dendritic cells, and HSPCs in KD patients before treatment

We further compared the expression features of major immune cell types and HSPCs between febrile patients/KD patients after treatment and KD patients before treatment. Most expression features among major immune cell types are similar to the overall expression features (Supplemental Table 1 and 2). The genes related to B cell receptor signaling pathway (IGHC3, IGCHM, IGHG4, IGHG1, IGLC3, GO:0050853) are also downregulated in B cells and plasma blast cells in KD patients before treatment (Figure 3A and 3B). It is noticed that the genes related to the biological processes of acetylation, cell cycle and cell division (UBE2C, HSPD1, HSPE1, MT2A, MYC, GO: KW-0007, KW-0131 and KW-0132) are selectively upregulated in DCs and HSPCs in KD patients before treatment (Figure 3C and 3D). We further compared the HSPCs expression profiles between febrile control and KD patients before treatment, since HPSCs give the rise to the other types of cells. There are 122 upregulated genes in HSPCs in our dataset and 81 upregulated genes in HSPCs in GSE168732 dataset (Figure 4A). 14 HSPCs upregulated genes (FOS, BLVRB, MCM7, RASD1, HIST1H3H, SAT1, JCHAIN, NKG7, HSPA1A, SELENOS, S100A9, S100A8, FOSB, TYROBP) are shared by both datasets. They are mainly involved in the KEGG pathway of IL-17 signaling pathway (GO: hsa04657, FDR value = 0.008). GO analyses show that there are common cellular components (GO:0070062∼extracellular exosome, GO:0005576∼extracellular region, GO: KW-0964∼Secreted) are shared by KD before treatment from our dataset and GSE168732 dataset, although the genes underlying these cellular components are various in two datasets (Figure 4B and 4C). Interestingly, the biological process of immune response (GO:0006955) is upregulated in the HSPCs of our dataset while the biological process of inflammatory response (GO:0006954) is upregulated in the HSPCs of GSE168732 dataset. Both immune response and inflammatory response are an innate function of immune cells, which proposes a premature immune activation propensity in the HSPCs of KD.

**Figure 3.**
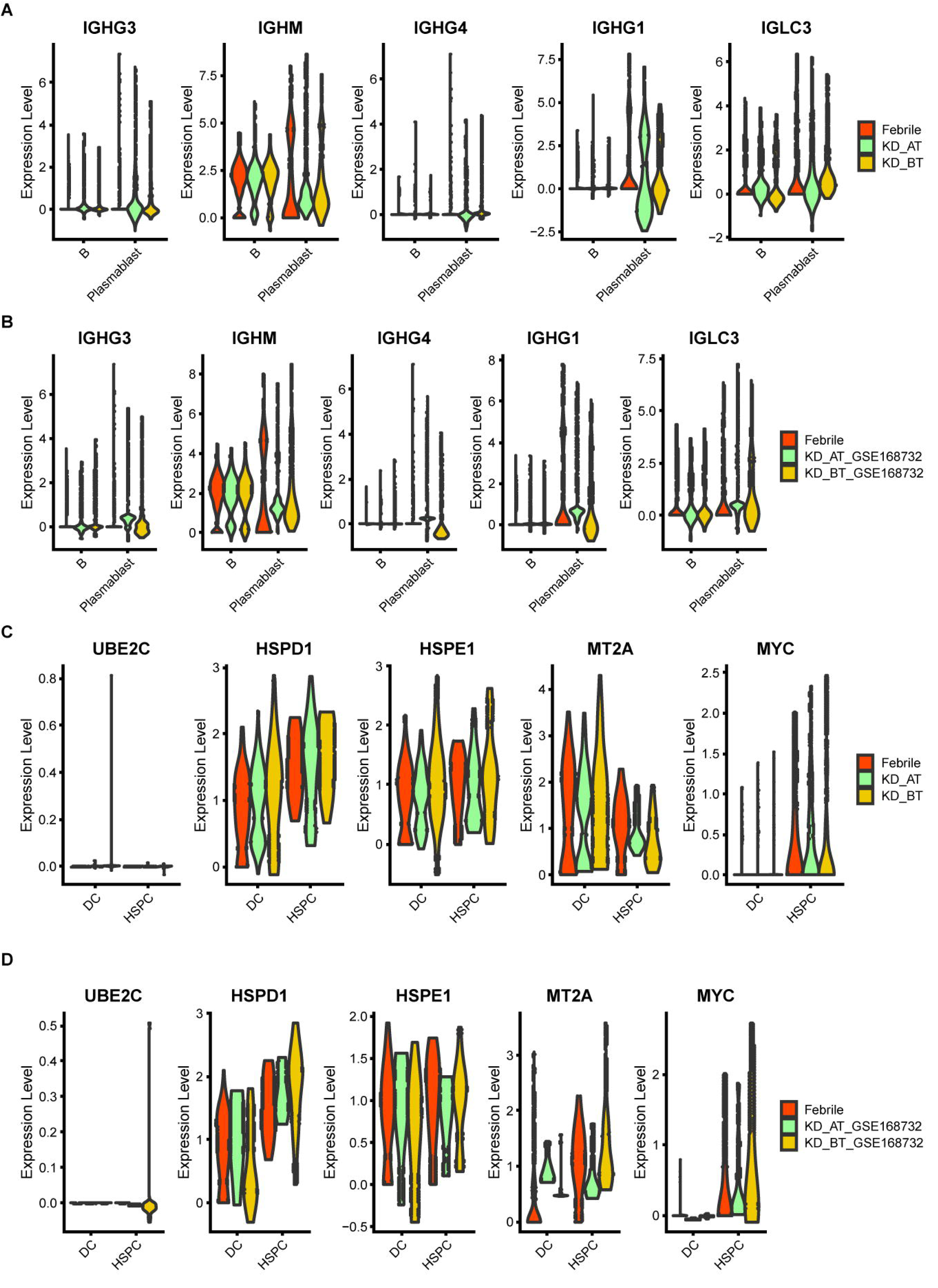
Violin plot for B cell signaling pathway genes and cycle cell genes in our dataset and GSE168732 dataset. A. The expression level of B cell signaling pathway genes in B cells and plasma blast cells of our dataset. B. The expression level of B cell signaling pathway genes in B cells and plasma blast cells of GSE168732 dataset. C. The expression level of acetylation, cell cycle and cell division genes in HSPCs and DC cells of our dataset. D. The expression level of acetylation, cell cycle and cell division genes in HSPCs and DC cells of GSE168732 dataset.

**Figure 4.**
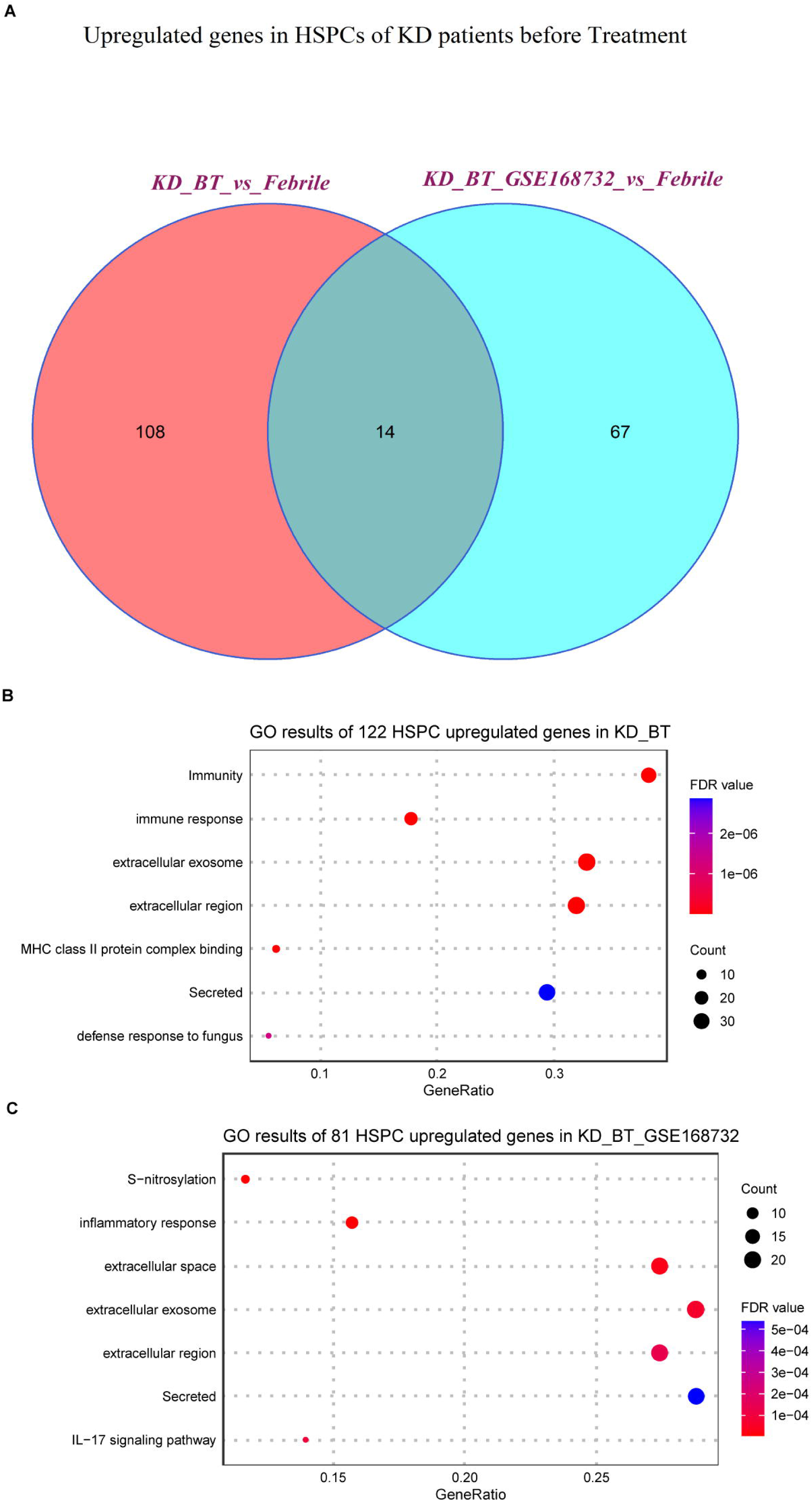
Expression analyses of HSPCs in our dataset and GSE168732 dataset. A. Venn diagram of upregulated genes in HSPCs for KD patients before treatment in our dataset and GSE168732 dataset. B. GO term enrichment analysis of 122 upregulated genes in HSPCs for KD patients before treatment of our dataset. C. GO term enrichment analysis of 81 upregulated genes in HSPCs for KD patients before treatment of GSE168732 dataset.

### Single-cell trajectory analyses of PBMCs in febrile and KD patients before/after treatment

UBE2C, HSPD1, HSPE1, MT2A, and MYC are the genes known to participate in the biological processes of cell cycle and cell division (GO: KW-0007, KW-0131 and KW-0132). Their abnormal upregulation in the HSPCs of KD before treatment could influence their descendant cells. We used pseudo-time analysis to reconstruct cell developmental trajectory of PBMCs in febrile and KD patients before/after treatment. Since HSPCs are the starting point of blood cell development, they are used to root the cell developmental trajectory of each dataset. The pseudo-time analyses show that the PBMCs in febrile and KD patients after treatment can be divided into five states while there are only three states of PBMCs in KD patients before treatment (Figure 5A, 5B, 5C 5D and 5E). It proposes that KD patients before treatment have a shorten cell development route and an ill cell differentiation outcome. The number of DEGs genes in cell developmental trajectory detected by Monocle 2 package are 15364, 14819, 15827, 12819 and 11867 in febrile patients, KD patients before treatment, KD patients after treatment, KD patients before treatment of GSE168732, and KD patients after treatment of GSE168732, respectively. Human PBMCs are known to form different cell lineages. We use a set of classic PBMC cell markers to identify the cell lineage for each state (Figure 5F and Supplemental Figure 2B) (24). For febrile patients, five states of their PBMCs are myeloid lineage (state 1), lymphoid lineage 1 (state 2), mixed lymphoid lineage 1 and 2 (state 3), erythro-megakaryocytic lineage (state 4), and lymphoid lineage 2 (state 5) (Supplemental Figure 2a to 2e). For KD patients before treatment, three states of their PBMCs are mixed myeloid and erythro-megakaryocytic lineages (state 1), myeloid lineage (state 2), and mixed lymphoid lineage 1 and 2 (state 3) (Supplemental Figure 3A to 3D). For KD patients after treatment, five states of their PBMCs are mixed myeloid and erythro-megakaryocytic lineages (state 1), lymphoid lineage 1 (state 2), mixed lymphoid lineage 1 and 2 (state 3), mixed myeloid lineage and lymphoid lineage 2 (state 4), and mixed lymphoid lineage 1 and 2 (state 5) (Supplemental Figure 4A to 4F). For KD patients before treatment of GSE168732, three states are lymphoid lineage 1 (state 1), mixed myeloid and erythro-megakaryocytic lineages (state 2), and mixed lymphoid lineage 1 and 2 (state 3) (Supplemental Figure 5A to 5D). For KD patients after treatment of GSE168732, mixed lymphoid lineage 1 and 2 (state 1), mixed lymphoid lineage 1 and 2 (state 2), lymphoid lineage 1 (state 3), mixed lymphoid lineage 1 and erythro-megakaryocytic lineage (state 4), and mixed lymphoid lineage 1 and myeloid lineage (state 5) (Supplemental Figure 6A to 6F). The cell lineage analyses show that only febrile patients have a clear cell lineage division while KD patients before/after treatment have more than one mixed lineage. Compared with KD patients before treatment, KD patients after treatment have more B cell lineages (lymphoid lineage 1). These results propose that B cell developmental dysfunction is the potential etiology of KD at cellular level.

**Figure 5.**
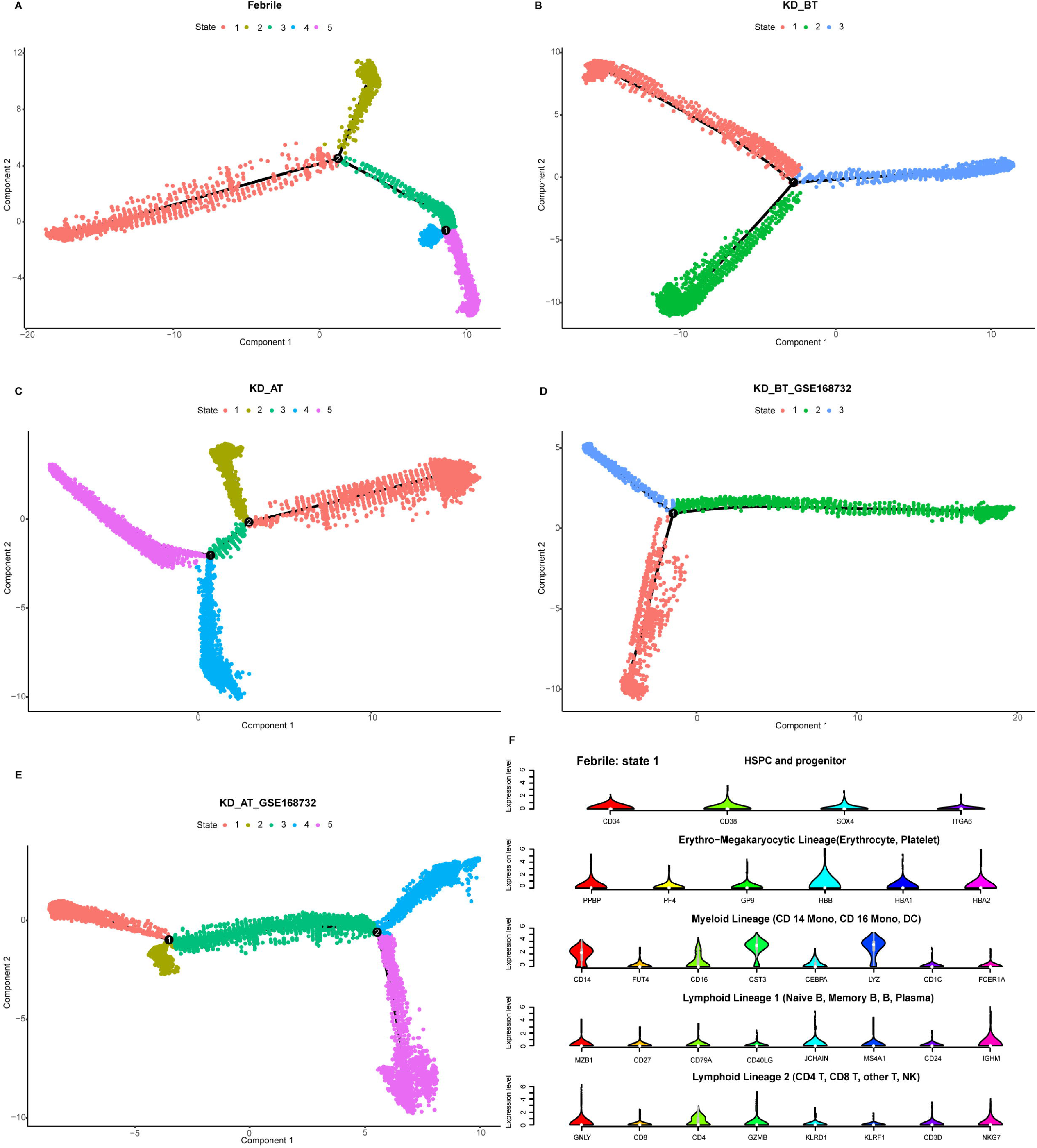
Pseudo-time analysis of cell developmental trajectory in our dataset and GSE168732 dataset. A. The differentiation trajectory of all cells in febrile patients by states in our dataset. B. The differentiation trajectory of all cells in KD patients before treatment by states in our dataset. C. The differentiation trajectory of all cells in KD patients after treatment by states in our dataset. D. The differentiation trajectory of all cells in KD patients before treatment by states in GSE168732 dataset. F. The differentiation trajectory of all cells in KD patients after treatment by states in GSE168732 dataset. F. The canonical markers of five cell lineages for state 1 in febrile patients in our dataset.

**Figure 6.**
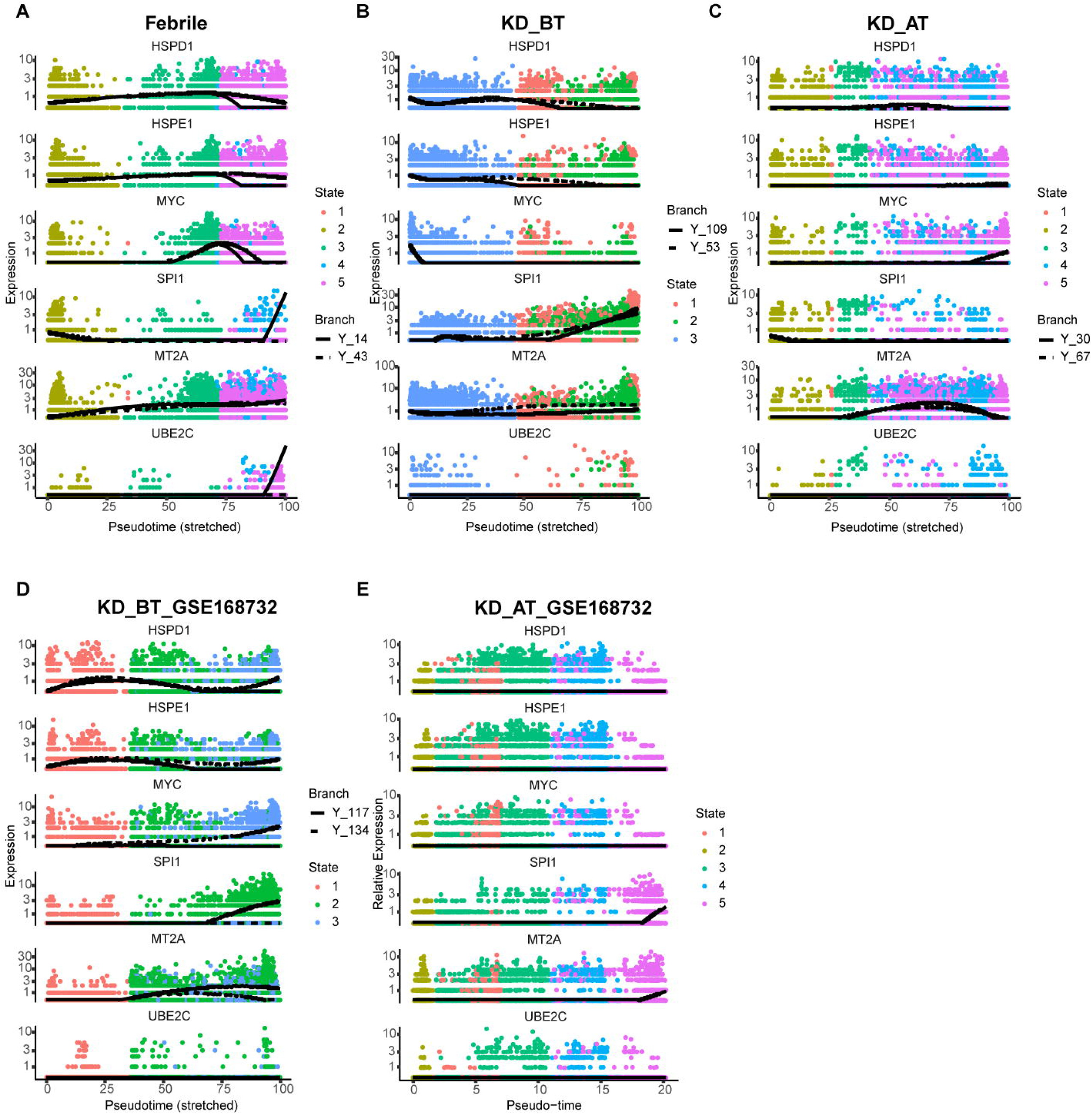
Pseudo-time analyses of expression dynamics of six cell-cycle related genes in our dataset and GSE168732 dataset. A. Expression dynamics of six cell-cycle related genes in febrile patients of our dataset. B. Expression dynamics of six cell-cycle related genes in KD patients before treatment of our dataset. C. Expression dynamics of six cell-cycle related genes in KD patients after treatment of our dataset. D. Expression dynamics of six cell-cycle related genes in KD patients before treatment of GSE168732 dataset. F. Expression dynamics of six cell-cycle related genes in KD patients after treatment of GSE168732 dataset.

### Pseudo-time expression dynamic analyses of cell cycle related genes and SPI1 in febrile and KD patients before/after treatment

We have observed that the genes related to cell cycle and cell division are selectively upregulated in DCs and HSPCs in KD patients before treatment. Single-cell trajectory analyses further show that KD patients before treatment have a shorten cell development route and an ill cell differentiation outcome. Thus, we investigate the expression dynamics of these genes based on pseudo-time analysis. We also added SPI1 gene in pseudo-time expression dynamic analyses, because it is a key gene in controlling myeloid and B-lymphoid cell lineage development (25). The expression dynamics of UBE2C, HSPD1, HSPE1, MT2A, MYC, and SPI1 were plotted along cell developmental trajectory based on pseudo-time analysis. We found that all six genes demonstrate a different expression dynamic among febrile patients, KD patients before treatment, and KD patients after treatment (Figure 6A to 6E). HSPD1 and HSPE1 show an elevated expression level in the early cell development stage of KD patients before treatment in our dataset and GSE168732 dataset, which are both repressed in KD patients after treatment. In febrile patients, MYC has an expression peak in the middle-late stage of cell development and SPI1 is highly expressed in the early and late stages of cell development. In KD patients before/after treatment, the expression peak of MYC disappears in the middle-late stage of cell development and SPI1’s early expression is almost repressed. The expression of MT2A and UBE2C are repressed in KD patients before treatment. Interestingly, these six genes show a different expression dynamic between febrile patients and KD patients after treatment. Except MT2A, their expressions are repressed in KD patients after treatment compared with febrile patients. It indicates that IVIG treatment probably restores the B cell development dysregulation in KD patients through suppressing the expression of UBE2C, HSPD1, HSPE1, MYC, and SPI1.

### The developmental path of B cells in febrile patients and KD patients before/after treatment

The overall profiling of PBMCs in febrile patients and KD patients before/after treatment shows that febrile and KD patients after treatment have the larger number of plasma blast cells than KD patients before treatment in both our and GSE168732 datasets. The overall expression features also demonstrated that B cell activation pathway is repressed in KD patients before treatment. In order to elucidate the possible etiological mechanisms of KD, it is worth investigating the detailed developmental path of B cells in three sample datasets. Thus, we extracted all HSPCs and B cells from each sample dataset and used them to perform pseudo-time analysis. B cells could be divided into four subgroups including B naïve, B intermediate, B memory and plasma blast cells. The B cell developmental trajectories display seven states in febrile patients, three states in KD patients before treatment, seven states in KD patients after treatment, one state in KD patients before treatment of GSE168732, and three states in KD patients after treatment of GSE168732 (Figure 7A to 7E). This result indicates that the B cells are fully developed in febrile patients and KD patients after treatment than in KD patients before treatment. The different types of B cells were plotted along their developmental trajectories. The difference between the B cells in our dataset and GSE168732 dataset is that KD patients in our dataset have more states than KD patients in GSE168732 dataset. In both our and GSE168732 datasets, KD patients before treatment have a more shortened B cell development path than KD patients after treatment. It indicates that most B cells in KD patients before treatment might be in a transitional state compared with those in KD patients after treatment. CD24 is a marker gene usually expressed in transitional B cells (26, 27). We extracted B naïve, B intermediate, and B memory cells for five data samples and analyzed CD24 expression level in all of them. Its expression range is wider in KD patients before treatment in both our and GSE168732 datasets. Our further analysis shows CD24 is actually a cell maker gene for most B cells in KD patients before treatment, whose expression level is significantly higher in B naïve, B intermediate, and B memory cells compared with KD patients after treatment in our and GSE168732 datasets (Figure 7F). Furthermore, CD24 expression level is higher in our dataset than in GSE168732 dataset (Wilcox tests, p<0.05). It might explain the different number of states in these two datasets. In all five data samples, plasma blast cells are always at the end of B development path. It is consistent with the current knowledge of B cell development process. Thus, the insufficient number of plasma blast cells in KD patients before treatment is due to B cell developmental dysfunction whose main feature is the elevated expression of CD24. It proposes that B cells in KD patients before treatment are mostly arrested in a transitional state.

**Figure 7.**
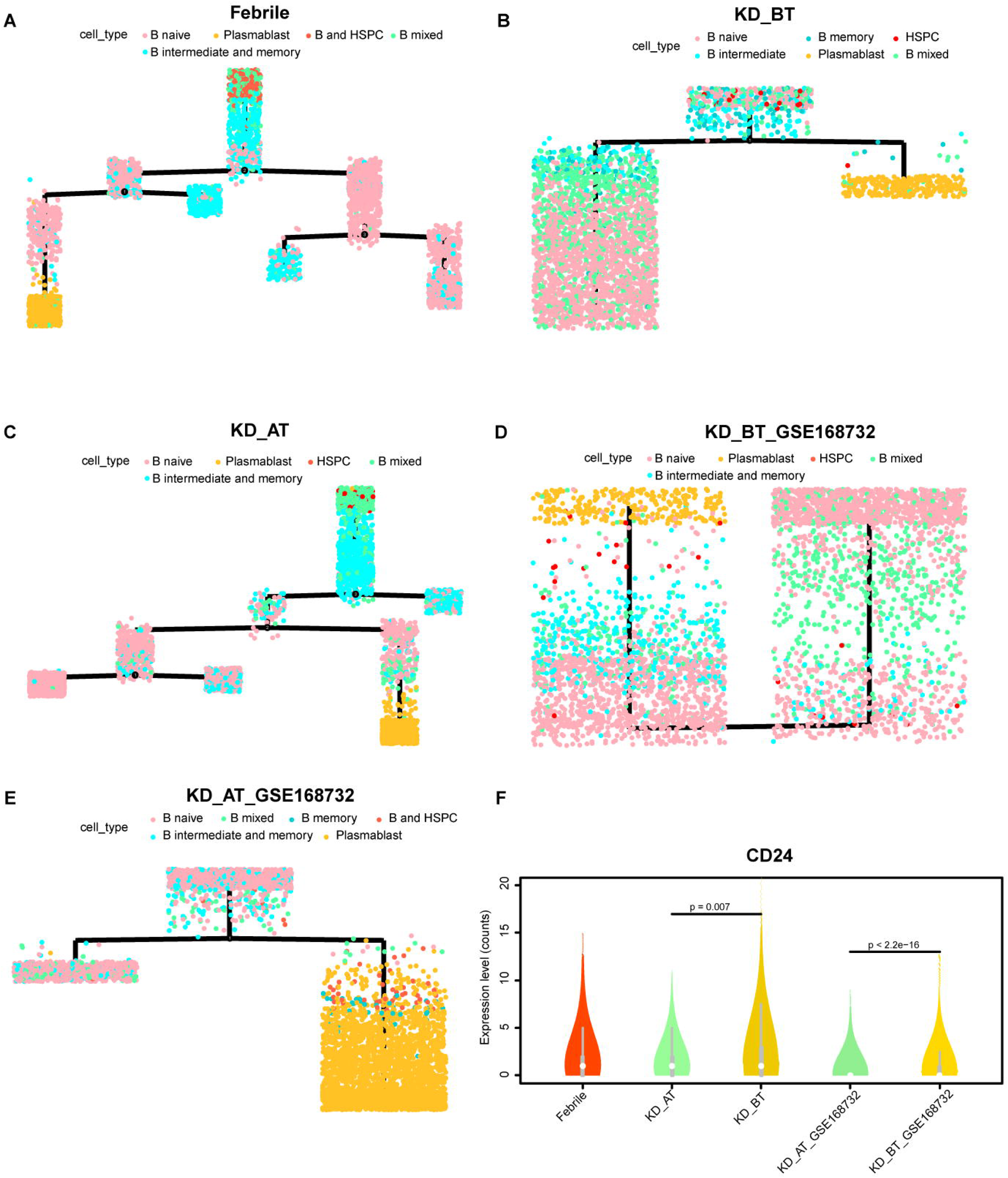
Pseudo-time analysis of cell developmental trajectory of HSPCs and B cells in our dataset and GSE168732 dataset. A. The differentiation trajectory of HSPCs and B cells in febrile patients by cell types in our dataset. B. The differentiation trajectory of HSPCs and B cells in KD patients before treatment by cell types in our dataset. C. The differentiation trajectory of HSPCs and B cells in KD patients after treatment by cell types in our dataset. D. The differentiation trajectory of HSPCs and B cells in KD patients before treatment by cell types in GSE168732 dataset. F. The differentiation trajectory of HSPCs and B cells in KD patients after treatment by cell types in GSE168732 dataset. F. The violin plot of CD24 expression in the B cells in our dataset and GSE168732 dataset.

## Discussion

KD was initially reported in 1967. Despite tremendous efforts, the etiology of KD remains unclear. Numerous etiology theories have been proposed for KD. The infection hypothesis suggests that an infectious agent leads to activation of the immune system in a genetically susceptible child who eventually developed KD (27). This theory is supported by the apparent seasonality of KD, the similar clinical features with other infectious diseases and the increased levels of inflammatory markers (28–30). However, none of these potential KD infectious agents have been confirmed. A large number of studies have revealed the significant activation of innate immune response and adaptive immune response in KD, which support the autoimmunity hypothesis. However, its low recurrence rate and the absence of a family history of autoimmune diseases in KD are inconsistent with the typical presentation of autoimmune disorders (1, 31, 32). In short, none of these theories of KD have been fully validated and they only partially account for the characteristics of KD. Until now, there is no consensus on its etiology.

Then hygiene hypothesis was introduced to explain the etiology of KD. Epidemiological and immunological data support it as a possible KD etiology and the development of B cell is a crucial factor in this hypothesis (13). In previous studies, the increase of B cell number in peripheral blood of acute KD patients has been documented. The infiltration of oligoclonal plasma cells producing IgA into the vascular wall of KD patients was also reported (33). A series of genome-wide studies identified several KD susceptibility genes are involved in B cell development and function (17–19). These findings have suggested that B cell related immunity is crucial in the development of KD. The overall single-cell expression features show that B cell receptor signaling pathway related genes are downregulated in KD before treatment while innate immunity response related genes are upregulated. These results confirm that B cell development is dysregulated in KD and such dysregulation could lead to excessive innate immune response. B cells play a key role in adaptive immunity. Their undergrowth would shift KD patients’ immune system to rely on innate immune response for self-defense (28). Unfortunately, such shift would lead to over-reaction of immune system and further invoke immune system to attack benign antigens. Thus, our findings are consistent with the concept of hygiene hypothesis. We further identified that HSPCs in our dataset and GSE168732 dataset overexpress the genes related to IL-17 signaling pathway which are mainly involved in inflammation. HSPCs in both datasets exhibit a premature immune activation propensity, which suggests that excessive immune response starts very early in KD patients and B cell development dysregulation could begin in bone marrow.

The analyses of expression features of major immune cell types in KD before treatment further reveal the possible genes behind B cell development dysregulation. First, the B cell receptor signaling pathway related genes are downregulated in B cells and plasma blast cells in KD before treatment. Their low expression level proposes that upstream activation signal is repressed for the B cells in KD patients is repressed and thus the B cells in KD patients have difficulties to differentiate into downstream cells such as plasma blast cells. Second, the selective upregulation of five cell cycle and cell division related genes in dendritic cells and HSPCs lead to produce a large number of premature B cells in KD before treatment. The untimely expression of cell cycle and cell division related genes generates the premature B cells which have poor expression of the B cell receptor signaling pathway related genes. They mutually contribute to the symptom of a large number of B cells and a small number of plasma blast cells observed in KD patients before treatment. The B cell developmental dysplasia in KD patients before treatment is also proven by clinical data which show that the IgG/M level is low in KD patients compared to febrile patients.

The pseudo-time analyses of KD before treatment in our and GSE168732 datasets has an impaired cell development trajectory compared with Febrile and KD after treatment. Canonical cell markers show Febrile patients have four clearly differentiated states which are myeloid lineage (state 1), lymphoid lineage 1 (state 2), erythro-megakaryocytic lineage (state 4), and lymphoid lineage 2 (state 5), but KD patients before treatment have mixed myeloid and erythro-megakaryocytic lineage and mixed lymphoid lineage 1 and lineage 2 (29). It suggests that B cell development dysregulation in KD actually have a deep root in the very early cell lineage differentiation which not only affect lymphoid lineage but also myeloid lineage. SPI1 is also known as hematopoietic transcription factor PU.1. It is a very important regulator gene in controlling gene expression during myeloid and B-lymphoid cell development (25). The gene expression dynamic along the cell development trajectory shows that SPI1 is highly expressed in the early and late stages of cell development in febrile patients but its early expression is repressed in KD before treatment. Its repression in KD before treatment is likely influenced by the early expression of HSPD1 and HSPE1, but how HSPD1 and HSPE1 repress the expression of SPI1 need to be studied in future. The overall expression feature analyses show that the hematopoietic cell lineage related genes are upregulated in KD before treatment. It further confirms that the SPI1 expression is repressed in KD before treatment, because it controls hematopoietic cells to differentiate into myeloid and lymphoid lineages. In hematopoiesis, HSPCs first give the birth to common myeloid progenitor cell and common lymphoid progenitor cell (29). The formers further split into myeloid lineage and erythro-megakaryocytic lineage and the latter further split into T lymphoid lineage (lymphoid lineage 1) and B lymphoid lineage (lymphoid lineage 2) (29). Thus, the repression of SPI1 expression in KD patients before treatment render them have only three developmental states in which myeloid lineage vs. erythro-megakaryocytic lineage and lymphoid lineage 1 vs. lymphoid lineage 2 are unable to separate from each other.

The examination of B cell developmental path further confirms KD before treatment have an ill B cell developmental trajectory as well. The B cells in KD before treatment has less states than those in febrile patients and KD after treatment. In both our dataset and GSE168732 dataset, IVIG treatment facilitates the B cells of KD to develop into more trajectory states and thus produce more plasma blast cells to perform normal adaptive immunity related functions. The expression dynamic analyses show that the expression of UBE2C, HSPD1, HSPE1, MYC, and SPI1 are repressed in KD after treatment. UBE2C, HSPD1 and HSPE1 participate in the biological process of cell cycle (GO: KW-0131) while MYC and SPI1 are two proto-oncogenes (30, 31). Their repression could stop cells from proliferating and thus render B cells unable to enter to next developmental fate. This speculation is supported by the high expression of CD24 in the B cells of KD before treatment which a marker for transitional B cells. The expression level of CD24 in the B cells of KD before treatment clearly demonstrates that they are arrested in a transitional state, but the CD24 expression is also downregulated after IVIG treatment. These results indicate that IVIG acts a general suppressive signal for KD patients’ immune system which would turn down its hyperactivity after receiving IVIG by facilitating the developmental fate of B cells.

We observed some variances between our dataset and GSE168732 dataset, especially in B cell developmental path analyses. Before treatment, the total B cells from the KD patients of our dataset have three states while the total B cells from the KD patients of GSE168732 dataset have only one state, which suggests that there exist some subtle pathological differences among different KD patients. There are about 10-20% of KD patients insensitive to IVIG treatment (32). These differences among KD population need to be investigated by future studies. Although whether lacking of common pathogens in living environment sets off KD still need to be investigated, our result unequivocally shows that KD has an immune system always ready for battle, i.e. a hyperactive immune state that could be easily activated by any antigen. Unfortunately, such hyperactive immune system is based on the disordered developmental fate of major immune cells, especially for B cells. There were several shortcomings in our study. Firstly, the sample size was limited, which affected the statistical power of this study. Secondly, there are very few public single-cell data for KD and we only found one available for this study. The future studies on KD’s pathological mechanisms requires more public data and endeavors from scientific community. Thirdly, this study was performed on PBMCs and might not reflect the local inflammatory responses developing in the coronary artery and the maturation process of hematopoietic cells in the bone marrow. Lastly, we only performed scRNA-seq in this study, which only reflected the single-cell transcriptome level of KD. Single-cell multi-omics sequencing can be considered to be applied in the further studies of KD.

## Materials and methods

### 1. Participants and statistical analysis

All participants were recruited from Shanghai Children’s Hospital. All manipulations were approved by the Ethics Committee of Shanghai Children’s Hospital (IRB number: 2022R121). Guardians of the participants had provided their informed consent in the study. KD was diagnosed according to the diagnosis criteria established by the American Heart Association (2). Two-dimensional echocardiography was used to examine whether there were coronary artery lesions during the acute and convalescent disease phases. Febrile patients were respiratory system infectious diseases admitted during the same period with KD patients.

Inclusion criteria for febrile group: (1) febrile patients (respiratory system infectious diseases) admitted during the same period with KD patients; (2) age matching with KD patients; (3) Fever duration matching with KD patients; (4) children who had not received steroids or immunosuppressive drugs within 14 days.

Exclusion criteria of all participants: 1, combined with autoimmune diseases; 2, combined with congenital malformations; 3, combined with metabolic diseases, etc.; 4, incomplete clinical data or inability to cooperate with the study. Peripheral blood samples collected from KD patients were on the 5th day after the onset of fever before IVIG treatment and 24h after IVIG treatment and subsidence of fever. Peripheral blood samples were also collected from three febrile patients immediately after admission.

Analysis was conducted using SPSS 27.0. Normally distributed continuous variables were expressed as mean ± standard deviation (s), with two independent samples t-test for between-group comparisons. Non-normally distributed measures were represented as median (M) (quartile 1 [Q1], quartile 3 [Q3]), and the Mann–Whitney U-test was used for the group comparisons. Count data were presented as the number of patients, using a four-compartment table χ^2^ test for between-group comparisons. P<0.05 was considered statistically significant.

### 2. Single-cell RNA sequencing and data analyses

#### 2.1 scRNA-seq library construction

Peripheral blood samples (2 mL each sample) were collected from the participants. PBMCs of participants were isolated according to standard density gradient centrifugation methods by using the Ficoll-Paque medium. The cell viability should exceed 90%. The single-cell library was constructed using the 5’ Library Kits. The cell suspension was loaded onto a chromium single-cell controller (10X Genomics) to generate single-cell gel beads in the emulsion (GEMs) according to the manufacturer’s protocol. Lysis and barcoded reverse transcription of polyadenylated mRNA from single cells was performed inside each GEM. Post-RT-GEMs were cleaned up, and cDNA was amplified. The barcoded sequencing libraries were generated using the Chromium Next GEM Single Cell V(D)J Reagent Kits v1.1 (10x Genomics) and were sequenced as 2 × 150-bp paired-end reads on an Illumina NovaSeq platform.

#### 2.2 Published data acquisition and selection

A set of published KD patients’ single-cell data was also used in our study to expand KD sample size (21). The published KD data could be retrieved from Gene Expression Omnibus (GEO) repository with the accession number of GSE168732 through NCBI website. Because we used three febrile patients as control in this study, three KD patients’ data were selected to get a paired dataset with our control from total six KD patients’ data of GSE168732. The selection procedure was as follows. First, we removed the low-quality samples from GSE168732. KD patient 1 before treatment of GSE168732 has a very low number of feature RNAs (average 700 feature RNAs) compared with the other five GSE168732 samples (average 1200 feature RNAs). KD patient 2 after treatment has a relative high percentage of expressed mitochondrion genes (about 10% of mitochondrion genes per cell) compared with the other five GSE168732 samples (about 7% of mitochondrion genes per cell). Thus, KD patient 1 and KD patient 2 were removed from this study for low quality. Second, we removed the sample with the excessive number of cells. KD patient 5 before treatment of GSE168732 has 8783 cells (vs. average 5585 cells in the other five before treatment datasets) and KD patient 5 after treatment of GSE168732 has 10201 cells (vs. average 8043 cells in the other five after treatment datasets). KD patients 5 of GSE168732 has about 25-50% more cells than the other five datasets of GSE168732 and is an outlier to be removed. Finally, KD patient 3, 4 and 6 of GSE168732 passed our selection criteria and were used in this study.

#### 2.3 scRNA-seq data processing

CellRanger (Version6.0.0) software was used to process the raw FASTQ files, align the sequencing reads to the GRCh38 reference transcriptome and generate a filtered UMI expression profile for each cell. Raw gene expression matrices were read into R (Version 4.2.1) and converted to Seurat objects. The number of genes, UMI counts and percentage of mitochondrial genes were examined to identify outliers. The following criteria was applied for quality control: total UMI count between 2,000 and 60,000, and mitochondrial gene percentage < 10%. After removal of low-quality cells, the count matrix was normalized by SCTransform method, which is based on a negative binomial regression model with regularized parameters. Then all datasets from the individual samples were integrated using the “FindIntegrationAnchors” and “IntegrateData” functions in Seurat (Version 4.1.1) (33). We identified “anchors” among all individual datasets with multiple canonical correlation analysis (CCA) and used these “anchors” to create a batch-corrected expression matrix of all cells, which allowed the cells from different datasets to be integrated and analyzed. The supervised principal component analysis (SPCA) was performed to reduction and the weighted nearest neighbor(wnn) graph-based clustering was used to identify cell clusters. The cell identities were determined with multimodal reference mapping in Seurat (Version 4.1.1) (33).

#### 2.4 Differential expression and functional enrichment analysis

DEG analysis for each cell type on the sample level following the recommendation of Bioconductor(34). The differential expression analysis was conducted between conditions by using Libra (Version 1.0.0) (35), which implements a total of 22 unique differential expression methods that can all be accessed from one function. We used “run_de” functions with pseudo bulk approach, implementing the DESeq2 (Version 1.36.0) (36) with a likelihood ratio test. GO analysis was performed using DAVID online resource (37). P-values were adjusted to FDRs. FDRs < 0.05 was chosen as the cut-off criterion indicating a statistically significant difference.

#### 2.4 Pseudo-time analysis of cell differentiation trajectories

Pseudo-time analysis of cell differentiation trajectories for each sample dataset was performed with R package Monocle 2 (38).The expression feature and inferred cell type for each sample data from Seurat result was used to construct the cell dataset for Monocle analysis pipeline. We used the Monocle built-in approach named “dpFeature” to detect the variable genes that define cell’s differentiation. Its advantages are needing no prior biological knowledge and discovering important ordering genes from data itself. Dimension reduction was performed with 2 max components and “DDRTree” method. HSPCs and B cells in each sample dataset for pseudo-time analysis were extracted according to the cell identities from Seurat result.

#### 2.5 Cell lineage verification

PBMCs are known to be differentiated into different lineages. The different cell lineage is verified with canonical cell markers in this study. The canonical cell markers were retrieved from CellMarker 2.0 database (http://bio-bigdata.hrbmu.edu.cn/CellMarker/) (24). We defined five cell lineages in this study which are HSPC and progenitor lineage with CD34, CD38, SOX4, and ITGA6 as marker genes, erythro−megakaryocytic lineage with PPBP, PF4, GP9, HBB, HBA1, and HBA2 as marker genes, myeloid lineage with CD14, FUT4, CD16, CST3, CEBPA, LYZ, CD1C, and FCER1A as marker genes, lymphoid lineage 1 (B cell lineage) with MZB1, CD27, CD79A, CD40LG, JCHAIN, MS4A1, CD24, and IGHM as marker genes, and lymphoid lineage 2 (T cell lineage) with GNLY, CD8, CD4, GZMB, KLRD1, KLRF1, CD3D, and NKG7 as marker genes. A custom R script based on vioplot library was developed to plot and verify all selected markers for each lineage.

## Data availability statement

The raw sequence data are available under restricted access because of data privacy laws, and access can be obtained by reasonable request to the corresponding authors.

## Author contributions

Z.W. and L.X. conceived and designed the project. L.X., Q.Q, S.S., L.C., X.J., W.L., and Q.L. collected samples and clinical information. G.D. and G.L. performed sequencing. L.S. and Q.L. performed bioinformatic analyses and discussed and interpreted the data. L.X., L.S, and Q.L. wrote the manuscript.

## Consent for publication

The author consented the right for publication.

## Acknowledgements

This work is jointly supported by National Natural Science Foundation of China (82170518) and the Shanghai Science and Technology Committee research Funding (22Y11909700). We would like to thank all the donors who contributed samples.

## Supplemental figure legends

**Supplemental figure 1.** Integrated single-cell profiling of PBMCs for our dataset and GSE168732 dataset. A. Integrated single-cell profiling of PBMCs for our dataset. B. Expression of canonical gene markers for each cell type in our dataset based on integration analysis. C. Integrated single-cell profiling of PBMCs for GSE168732 dataset. D. Expression of canonical gene markers for each cell type in GSE168732 dataset based on integration analysis. The inferred cell types are marked with different colors.

**Supplemental figure 2.** Pseudo-time analysis of cell developmental trajectory in febrile patients of our dataset. A. The differentiation trajectory of all cells in febrile patients by cell types in our dataset. B. The canonical markers of five cell lineages for state 1 in febrile patients in our dataset. C. The canonical markers of five cell lineages for state 2 in febrile patients in our dataset. D. The canonical markers of five cell lineages for state 3 in febrile patients in our dataset. E. The canonical markers of five cell lineages for state 4 in febrile patients in our dataset. F. The canonical markers of five cell lineages for state 5 in febrile patients in our dataset.

**Supplemental figure 3.** Pseudo-time analysis of cell developmental trajectory in KD patients before treatment of our dataset. A. The differentiation trajectory of all cells in KD patients before treatment by cell types in our dataset. B. The canonical markers of five cell lineages for state 1 in KD patients **before** treatment in our dataset. C. The canonical markers of five cell lineages for state 2 in KD patients before treatment in our dataset. D. The canonical markers of five cell lineages for state 3 in KD patients before treatment in our dataset.

**Supplemental figure 4.** Pseudo-time analysis of cell developmental **trajectory** in KD patients after treatment of our dataset. A. The differentiation trajectory of all cells in KD patients after treatment by cell types in our dataset. B. The canonical markers of five cell lineages for state 1 in KD patients after treatment in our dataset. C. The canonical markers of five cell lineages for state 2 in KD patients after treatment in our dataset. D. The canonical markers of five cell lineages for state 3 in KD patients after treatment in our dataset. E. The canonical markers of five cell lineages for state 4 in KD patients after treatment in our dataset. F. The canonical markers of five cell lineages for state 5 in KD patients after treatment in our dataset.

**Supplemental figure 5.** Pseudo-time analysis of cell developmental trajectory in KD patients before treatment of GSE168732 dataset. A. The differentiation trajectory of all cells in KD patients before treatment by cell types in GSE168732 dataset. B. The canonical markers of five cell lineages for state 1 in KD patients before treatment in GSE168732 dataset. C. The canonical markers of five cell lineages for state 2 in KD patients before **treatment** in GSE168732 dataset. D. The canonical markers of five cell lineages for state 3 in KD patients before treatment in GSE168732 dataset.

**Supplemental figure 6.** Pseudo-time analysis of cell developmental trajectory in KD patients after treatment of GSE168732 dataset. A. The differentiation trajectory of all cells in KD patients after treatment by cell types in GSE168732 dataset. B. The canonical markers of five cell lineages for state 1 in KD patients after treatment in GSE168732 dataset. C. The canonical markers of five cell lineages for state 2 in KD patients after treatment in **GSE168732** dataset. D. The canonical markers of five cell lineages for state 3 in KD patients after treatment in GSE168732 dataset. E. The canonical markers of five cell lineages for state 4 in KD patients after treatment in GSE168732 dataset. F. The canonical markers of five cell lineages for state 5 in KD patients after treatment in GSE168732 dataset.

