## Supplementary figures and images for "The Mechanism underlying B cell Developmental Dysfunction in Kawasaki Disease Based on Single-cell Transcriptomic Sequencing"

### Supplemental Figure1

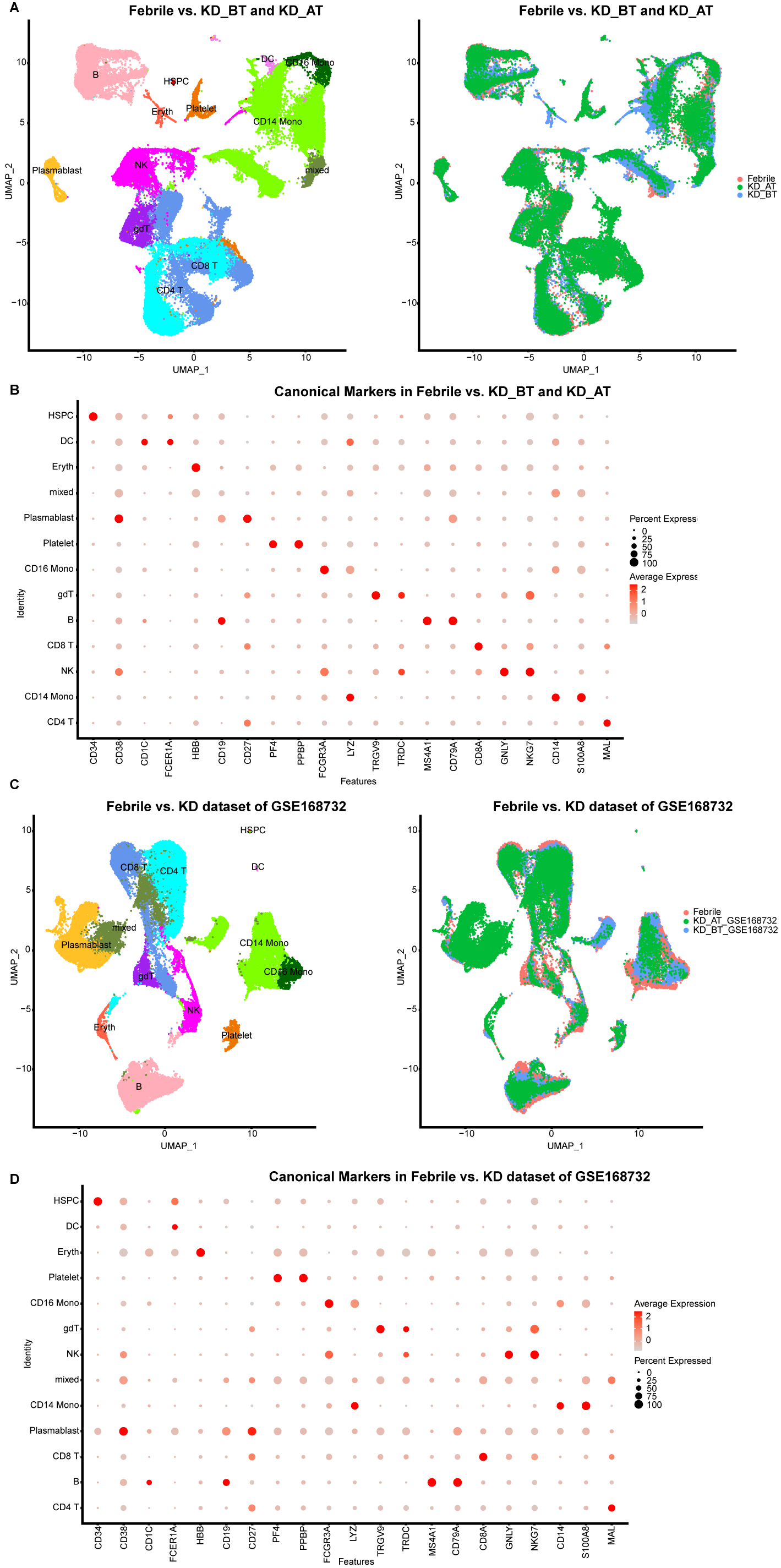

### Supplemental Figure2

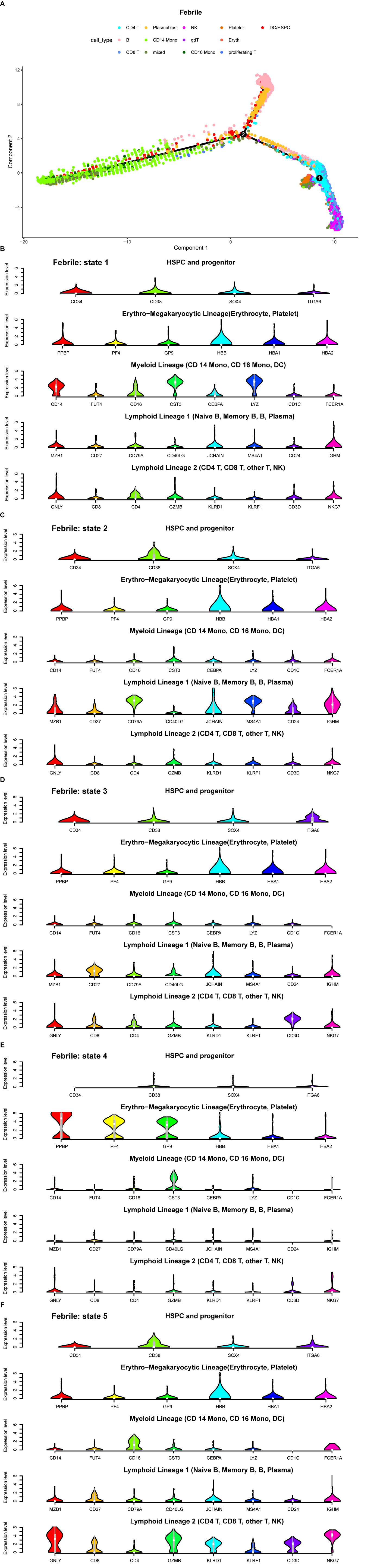

### Supplemental Figure3

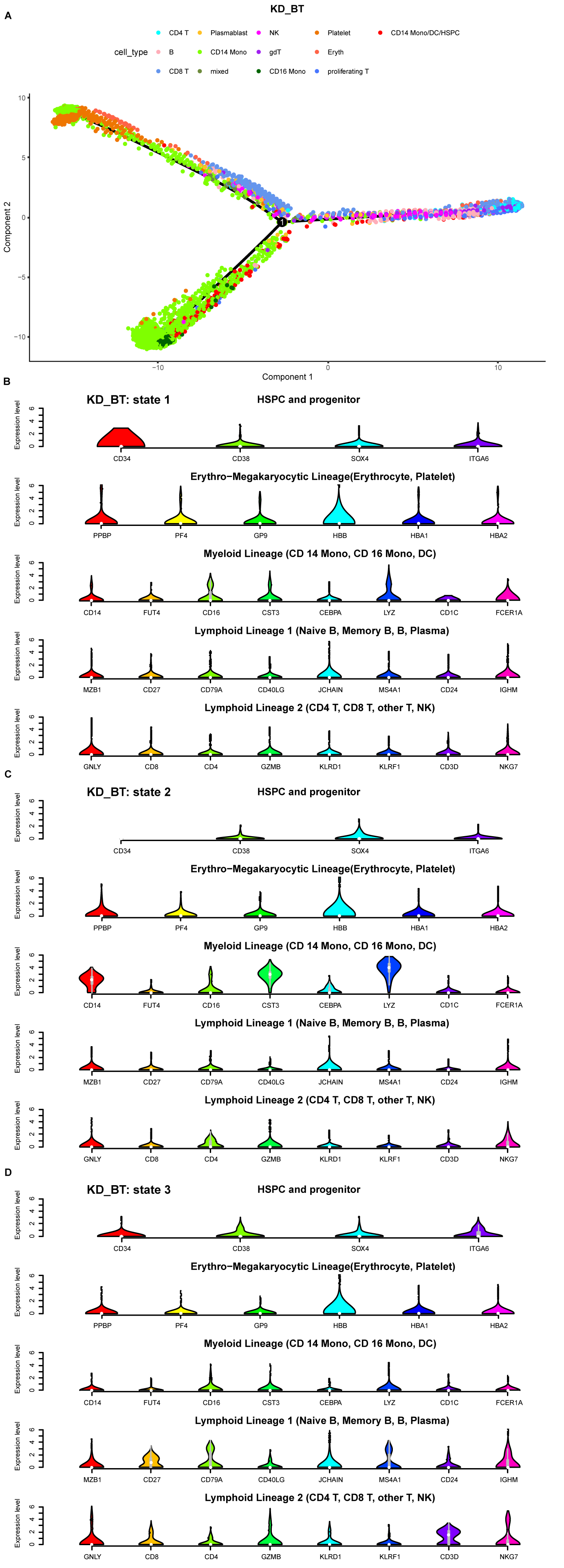

### Supplemental Figure4

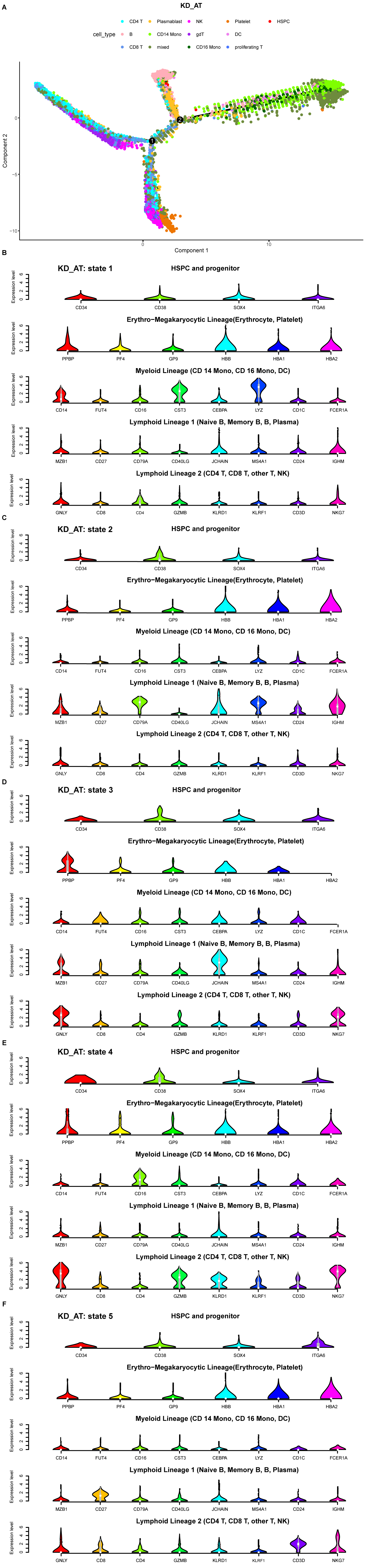

### Supplemental Figure5

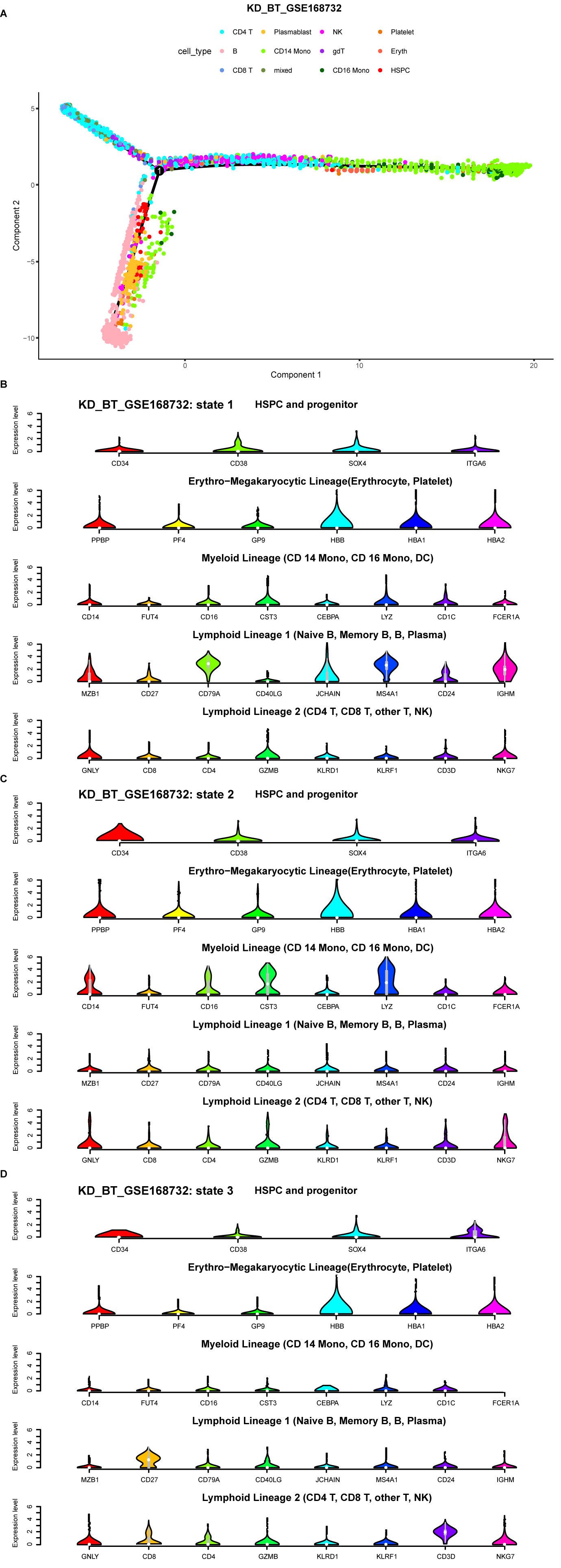

### Supplemental Figure6

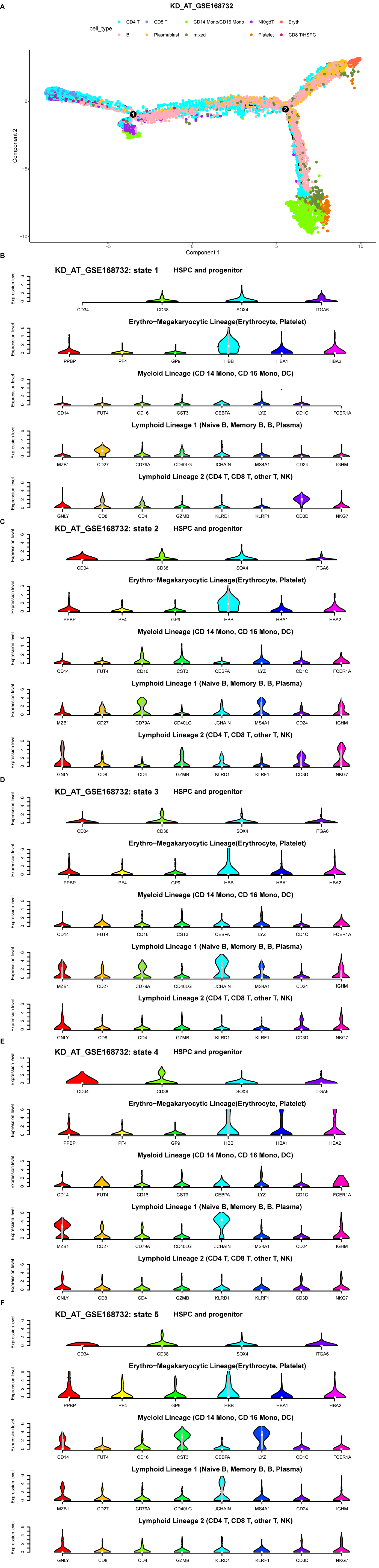
